# Energy balance; choices between energy spend on migration and mating influences the life cycle of the pond bat (*<u>M</u>yotis dasycneme*)

**DOI:** 10.1101/645309

**Authors:** Anne-Jifke Haarsma, Peter H.C. Lina, Aldo M. Voûte, Gerhard H. Glas, Henk Siepel

## Abstract

During autumn in the temperate zone, insectivorous male bats face a profound energetic challenge, as in the same period they have to make energy choices related to hibernation, mating and migration. We found evidence by looking at biometric measurements that male pond bats (*Myotis dasycneme*) are pre-occupied with mating and lose weight, while simultaneously females are accumulating fat. Our purpose was to characterize the known hibernacula in terms of male or female bias, and subsequently compare their population trend during two study periods, between 1930-1980 and 1980-2015. Our findings include evidence of colonisation of winter roosts in formerly unoccupied areas and consequently a change in the migration patterns of the male population of this species. As male bats do not assist to raise their offspring, males have abundant time to restore their energy balance after hibernation. Our results suggest that choosing a hibernacula closer to the summer habitat not only decreases energy cost needed for migration, it also lengthens the mating season and presumably also has the additional advantage of increased paternity. Additionally, these findings have important conservation implications, as male and female biased hibernation assemblages may differ critically in terms of microclimate preferences.

## Introduction

In contrast to bird migration, bat migration and its specifics are virtually unknown, such as its relation to summer distribution patterns, or how bats cope with energetic challenges and changing habitat. Bat species in the Northern hemisphere are almost all insectivorous., Due to lack of food resources, winters are spend in hibernation. In most species, male and female bats live in segregated summer habitats [1, 2; 3, 4, 5]. In autumn sexes meet in mating roosts, the male roosts, or the winter roosts [6, 7, 8]. The migration behaviour of bats between their summer and winter roosts is unique. Birds and large ungulates move from areas of low or decreasing resources (i.e. food) to areas of high or increasing resources [9, 10]. In contrast, the direction of migration in bats is mostly determined by the location of hibernacula with the suitable physiological requirements [11]. As a consequence, bats in the Northern hemisphere are sometimes observed migrating in northern directions from their summer roosts [12]. Among bats, distances between summer and winter habitat of more than several hundreds of kilometres are relatively uncommon. Exceptions in Europe are, for example, *Nyctalus noctula* and *Pipistrellus nathusii*, with the longest observed distances of up to 3,000 km [12]. Such distances are still meagre compared to the tens of thousands of kilometres some birds of similar sizes sometimes undertake [13]. Nonetheless bat migration has a substantial impact on energy balance [14, 15].

There are profound differences between the energy choices related to hibernacula selection for male and female bats [6]. Migration prior to hibernation not only takes a considerable energy cost, it also involves a long time spend with flying, perhaps alternated with fuel accumulation and time spend waiting for optimal conditions. Migration season and mating season overlap, therefore time and energy spend on migration is not available for mating. Although applicable for both male and females, this trade-off is probably more important to males, as lacking in mating activities combined with mating in suboptimal time and place may decrease their chances of fathering offspring. As male bats do not assist in to raise their offspring, males have abundant time to restore their energy balance after hibernation. In contrast, female bats begin their gestation in March/ April and are under considerable time pressure to return to their maternity roost and raise their offspring [16]. Males, however, are thought to spend more energy in mating before hibernation, which puts them for the dilemma of mating or migrating. Papers which present interesting life history trade-offs are plentiful, most are about female choice, age of first reproduction, in combination with longevity and hibernation choices [e.g. 17, 18]. Several studies have shown sex influenced differences in distribution patterns [3, 19], indicating that reproductive and energetic choices may also play a role in migration behaviour. In this paper we present an unique example of how choices of the males affect life history.

Here we characterize the known hibernacula of the pond bat, *Myotis dasycneme*, in terms of male or female bias, and subsequently compare their population trend during two study periods, between 1930-1980 and 1980-2015. To do so, we first imputed the missing values in the dataset. We predicted energy spend during mating cannot be spend on hibernation. Hence, we assessed the changes of the body condition index of bats during summer period. We compared migration distance, calculated by using mark recapture data, in the period before and after the settlement in the new hibernacula. Based on previous research [20], we hypothesized that the mating and hibernation are trade-offs, and thus we expected that a change in mate site location will lead to a change in migration distance.

## Methods

### Study area, periods and dataset

The study area covered the whole of the Netherlands, Belgium and East Frisia (northwest Germany). This area is considered the habitat of the “West European pond bat population”, an isolated population at its western limit of its distribution range [based on 21]. Hibernacula of this population are found along the southern and eastern border of its range, in Belgium, North France, West Germany and the southern part of the Netherlands. Since 1980 hibernacula sites also include several bunkers in Zuid-Holland and Gelderland (the Netherlands). Maternity roosts of this population are predominantly found in the centre of its range, in the Dutch lowlands, with male roost along the edges of the female habitat [22].

We defined two study periods, data collected between 1900 and 1980 (‘the historical dataset’) and data between 1980 and 2015 (‘the recent dataset’). The historical data were gathered from literature [23, 24, 25, 26, 27, 28, 29] and natural history museum collections. The recent data were gathered from the Dutch Mammal Society and from own observations by the first author [22, 30, 31]. These data were supplemented by data gleaned from publications [23, 32, 33, 34, 35, 36] and atlases, such as the atlas of the bats of the Netherlands [27], the Flemish region of Belgium [37, 38] and for several atlases of German Federal States [39, 40, 41]. Records of marked bats with rings and their biometry data were gathered from the first author and from the former Laboratory for Animal Ecology and Taxonomy of Utrecht University, former office of third author.

### Ringing and biometry

All available mark and recovery data (ringing) of both the historical and recent migration research were digitized. Observations include location and date of capture, species, sex and ring number. The latest observations in the recent dataset also include biometric measurements (forearm length, body weight) and information about age and reproductive status [42]. The biometric measurements were used to derive the body condition index of each individual. Bat captures were carried out under license from the Dutch Ministry of Economic affairs, (permits FEF27b/2002/034, FF/75A/2003/150, FF/75a/2006/013, FF/75a/2008/033, FF75a2012/37a and with permissions of all site owners. Ringing was carried out under license of Animal Experiments Committee UDEC 02036, 06058, 07124, 98055 and Alt 09-01). All bats were released within one hour, at the point of capture.

### Winter hibernation surveys

Hibernation surveys are often included in national monitoring programs [43, 44]. Sites are visited once each winter systematically searched with torch light until all visible bats were found and identified (without disturbing the bats). Although winter monitoring has its biases [e.g. 45, 46, 47], the data can be used to estimate coarse trends in abundance. For this paper we selected winter roosts with three or more records of three of more pond bats in one or both of the study periods. Only data from sites with long term data series (from the hibernacula in the Dutch provinces of Zuid-Holland, Gelderland and Limburg) were used for analysis of abundance. Unfortunately, due to a change in regulations, legal access to the Dutch limestone mines needed to carry out monitoring surveys has become problematic since 2011. Hence, we present winter data in this particular area only up to 2011.

### Statistical analysis

We calculated trends and annual abundance using General Linear Models (GLMs) with a Poisson error distribution, implemented in the software TRIM [48]. This programme is able to derive the values of missing observations based on previous and succeeding values. We used the linear trend model with all change points included (this is because of the absence of observation in 1949). Non-significant change points were excluded stepwise. In this paper winter abundance data are expressed as imputed annual abundance and abundance indices (using the first year as a base year, i.e. 100%).

## Results

### Mating effects, weight of males during autumn

The biometric measurements of the marked pond bats, showed that male pond bats are on average a few percent smaller and lighter than females. Analysis of our dataset revealed that changes of the body condition index of both sexes do not occur simultaneously, each is related to its respective reproductive cycle (Fig. 1). Near week 34, after their offspring is fully weaned, females start accumulating fat reserves. The pond bat mating season starts in week 28, from this time males are preoccupied with mating activities and as a consequence lose weight. Instead of accumulating fat, the weight of males drops and they start falling behind. Just before the start of the winter, 4-8 weeks later than females, males reach their pre-hibernation weight. This difference between male and female body condition index just before hibernation stresses the potential gain males may have in adapting their migrating behaviour.

**Fig. 1:**
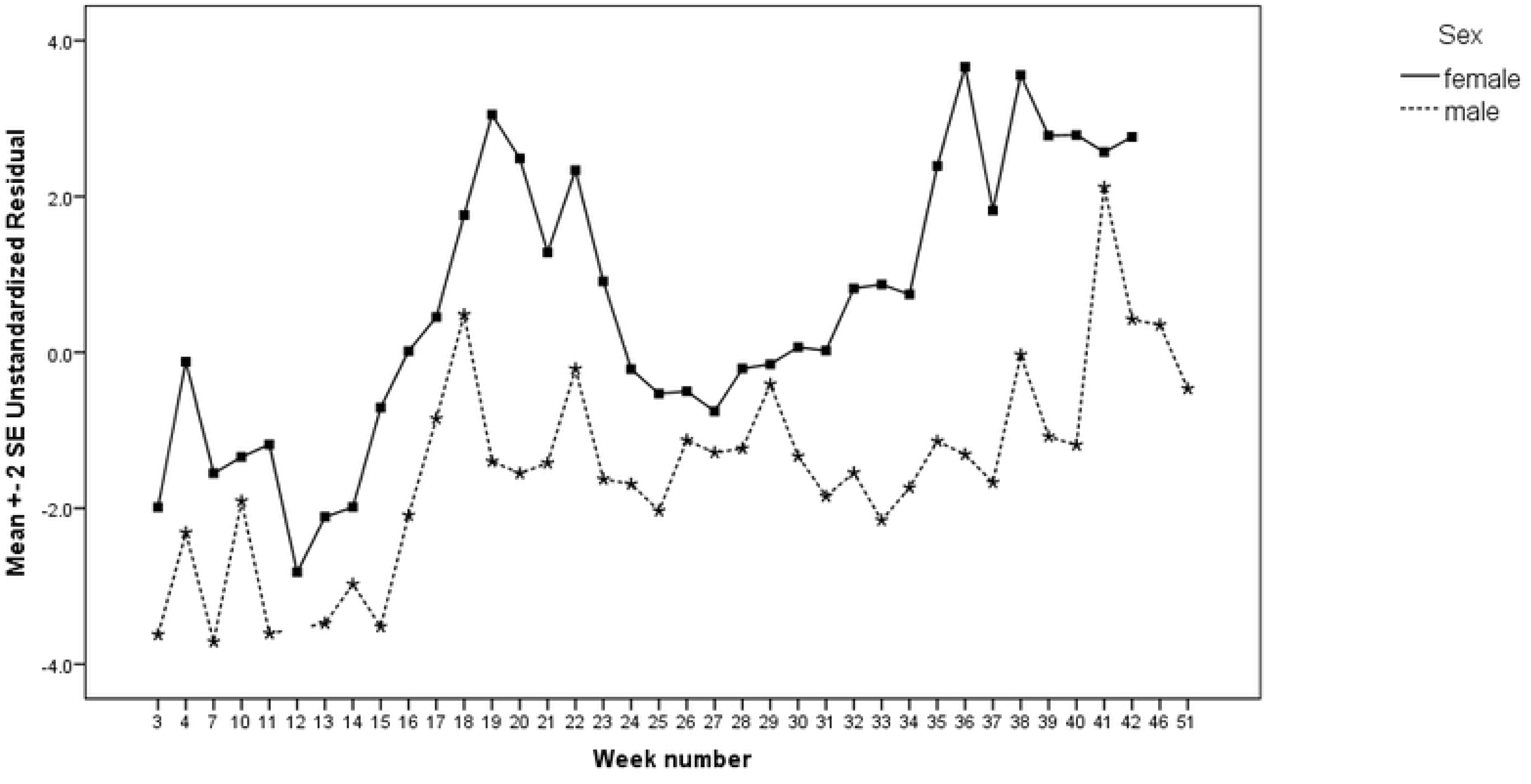
Fluctuations in the body condition index of the adult population during the year (data collected in period 2002-2015).

### Selection of new hibernacula near summer sites

A total of 113 unique hibernacula, 59 in Limburg, 38 in Zuid-Holland, 16 in Gelderland were surveyed during this study. The first colonization of hibernacula in formerly unoccupied areas occurred in the beginning of the 1980s (Fig. 2). A few pond bats (males) were found in 12 bunkers in the provinces of Zuid-Holland and Gelderland (hereafter referred to as the core bunkers), respectively along the western border and in the centre of the distribution range of the West European population. The bunkers were relative new sites; built during the Second World War (WOII) [e.g. 49]. Although sites adjacent to these core bunkers were visited annually, pond bats remained (nearly) absent until the year 1997 (referred to as the second colonization event). After an exponential increase in almost all sites (14 sites in Gelderland and 28 in Zuid-Holland), today’s numbers are 503 times higher than in the colonization year.

**Fig. 2:**
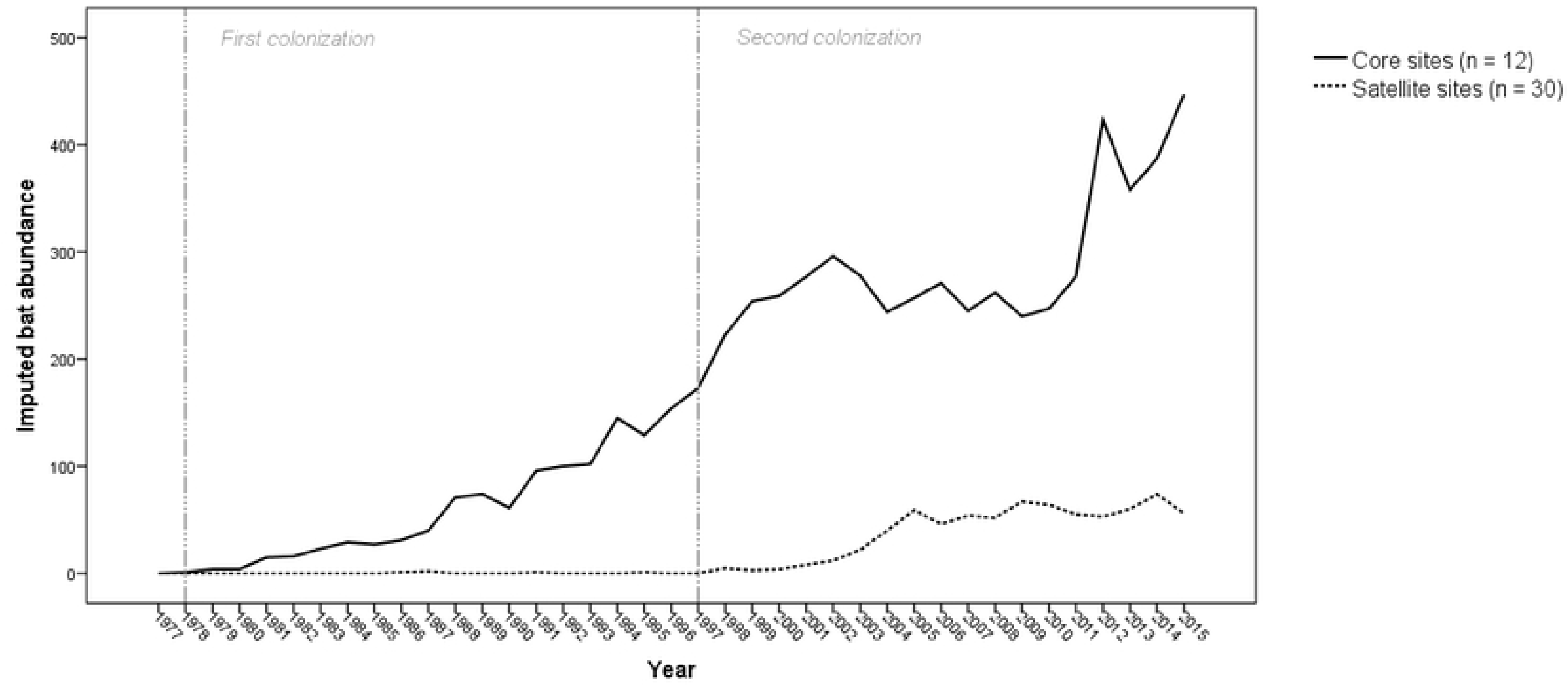
The (imputed) total abundance of pond bats in hibernacula in the provinces of Gelderland and Zuid-Holland after 1980. The two lines represent two types of hibernacula; core sites (12 roosts, unbroken line) and satellite sites (30 roosts, broken line).

This strong increase is in stark contrast to the population trend observed in Limburg and Southern Belgium. In a span of 20 years, between 1941 and 1961, the annual counts in 59 sites in the Netherlands Limburg decreased from 350 to 132 individuals. Instead of abandonment of sites, the average density of animals per site decreased from 6.1 to 3.2 pond bats. In the same period in southern Belgium the hibernation population decreased from an estimated 200 animals to a meagre 20 animals (11.6% decrease based on 58 hibernacula [50]. Since 1980, hibernation populations in Limburg and Southern Belgium remained more or less at the same reduced level after the decline.

### Assemblage in hibernacula is either male or female biased

The total western population of pond bats based on summer counts is estimated at approximately 24,000 bats (assumed females:males = 1:1) [21, 30]. A very small proportion of this population is observed during the winter, approximately 1,500 bats [21, 30]. Hibernation location of the rest of the population is unknown.

We presume (large) site-specific underrepresentation of the true numbers of hibernating bats is likely, due to variations of internal characteristics of each site (e.g. size, number of cracks and crevices) and the roosting ecology (pond bats are considered to hibernate mostly in cold crevices [51]. Research pioneers, such as Punt and Van Nieuwenhoven [52] and Kugelschafter [53], showed such an error can be considerable. Although our results might be biased based on the above statement, we made estimations of the male: female ratio in bunkers and limestone mines. We found that the bunkers are predominantly inhabited by males (approximately male:female = 8:1), while the limestone mines were predominantly female biased (before 1980 approximately male:female = 1:3 (based on original data gathered by Laboratory for Animal Ecology and Taxonomy of Utrecht University, after 1980 approximately male:female = 1:5).

### Change in migration distance

Before 1980 over 3,000 pond bats were captured and marked (Table 1a). A total of 1,214 pond bats have been ringed in their maternity roosts, marked animals were predominantly females and their offspring. An unknown number of bats were captured in their hibernacula in Southern Belgium, both Netherlands and Belgian Limburg and the Eifel region [54, 55, 23, 32, 28]. The reported recoveries of these ringed animals digitized by us showed that the animals spending the summer in the Netherlands migrated long distances to hibernacula in the Southern Netherlands (Limburg), Belgium (Namur) and in Germany (Fig. 3a). Animals from the Dutch province of Friesland were often recovered to the south in Southern Limburg in the Netherlands and to the east in Germany with a preference for the Teutoburg forest region [28, 56].

**Table 1a and b:**
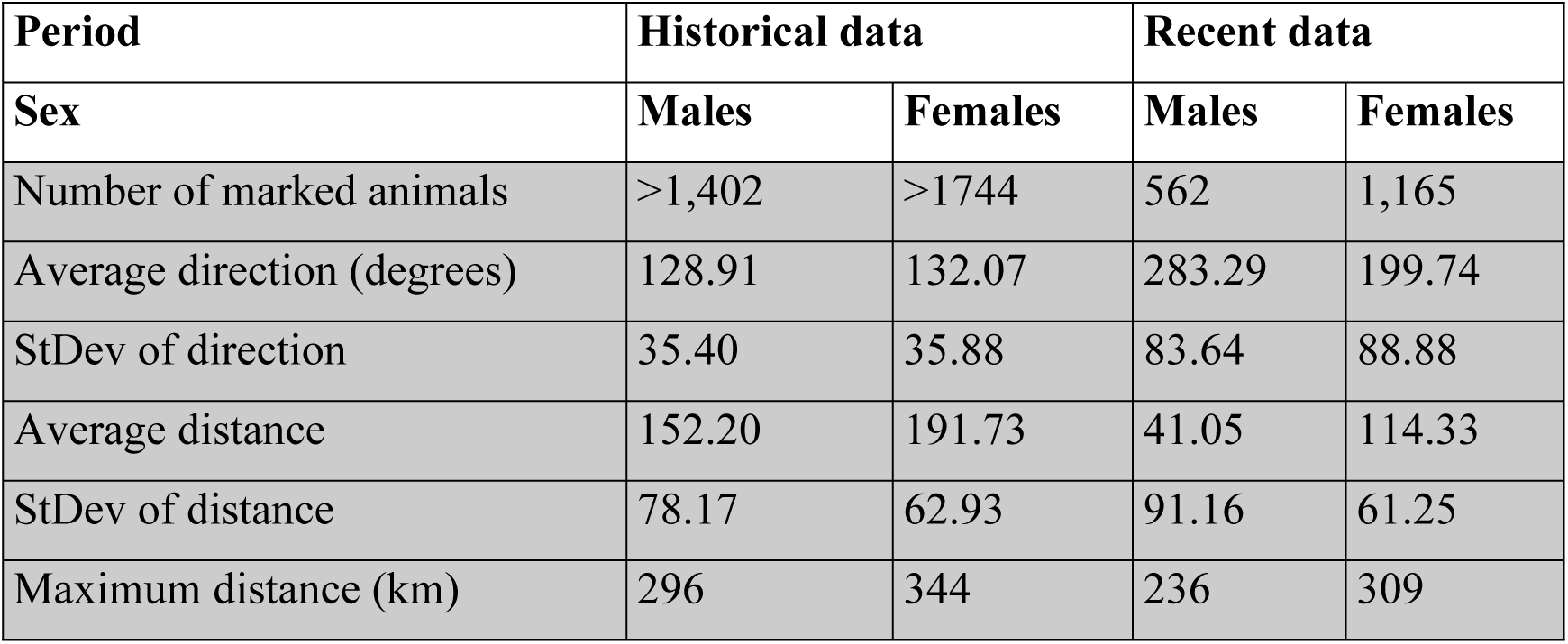
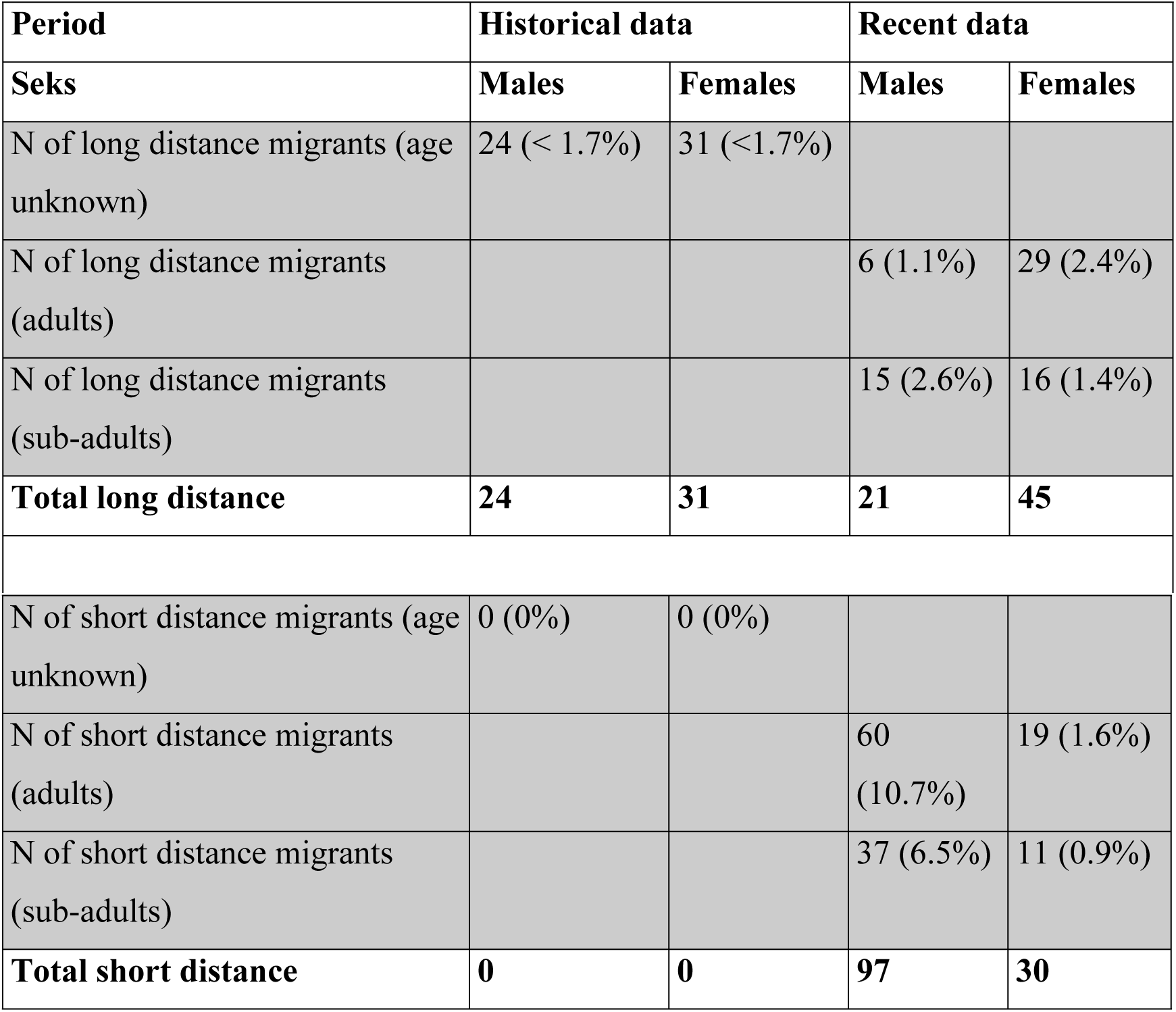
Comparison between migration behaviour based on historical data (1939 – 1973) and recent data (2002-2015). The numbers in parentheses are the percentage of the marked population, calculated for each gender. The direction of migration is the direction with the summer roosts as starting point (even if bats were originally captured during winter). Short distance are distances less than 50 km between the summer and winter roost, long distances are more than 50 km.

**Fig. 3a and b:**
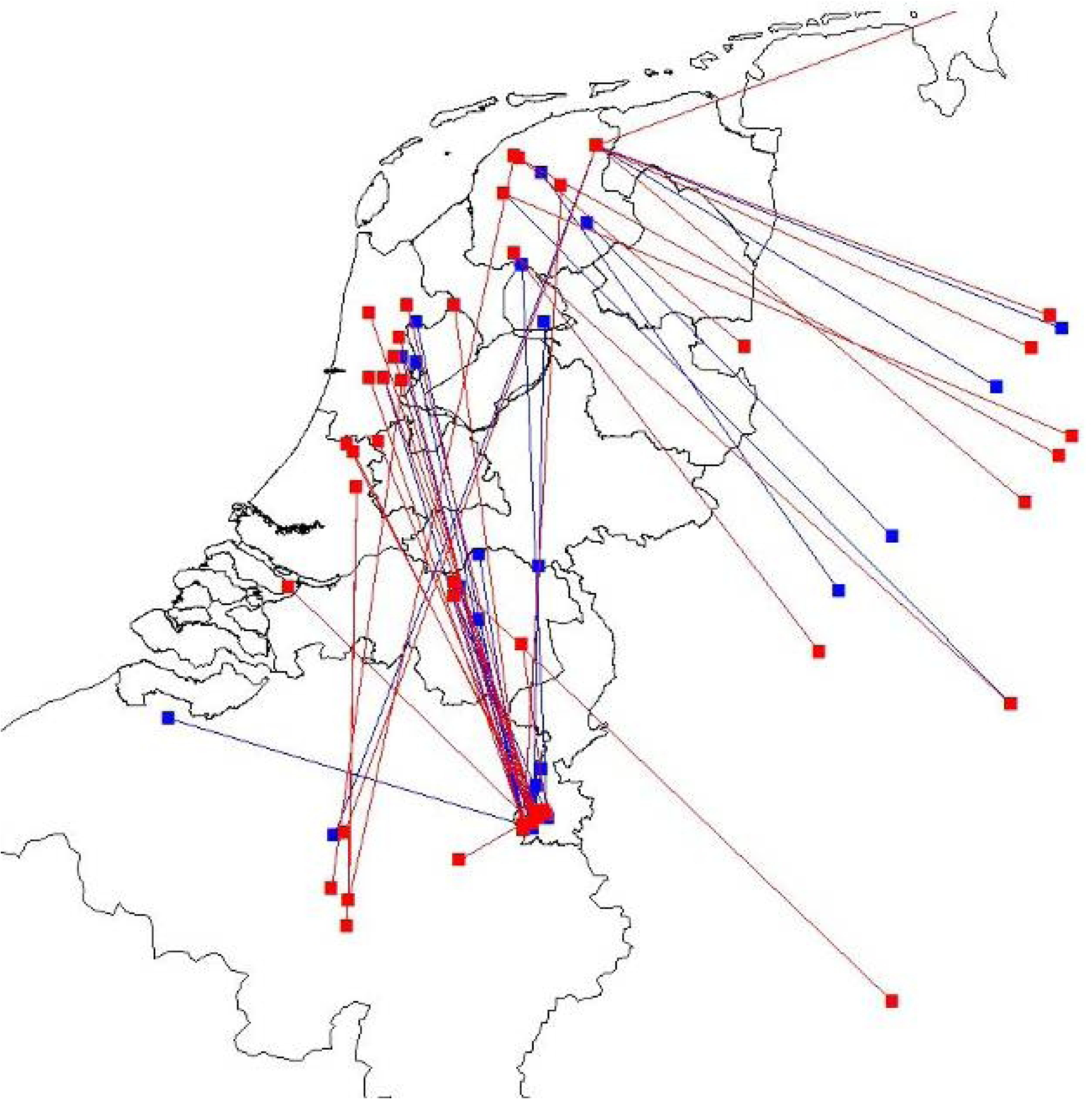
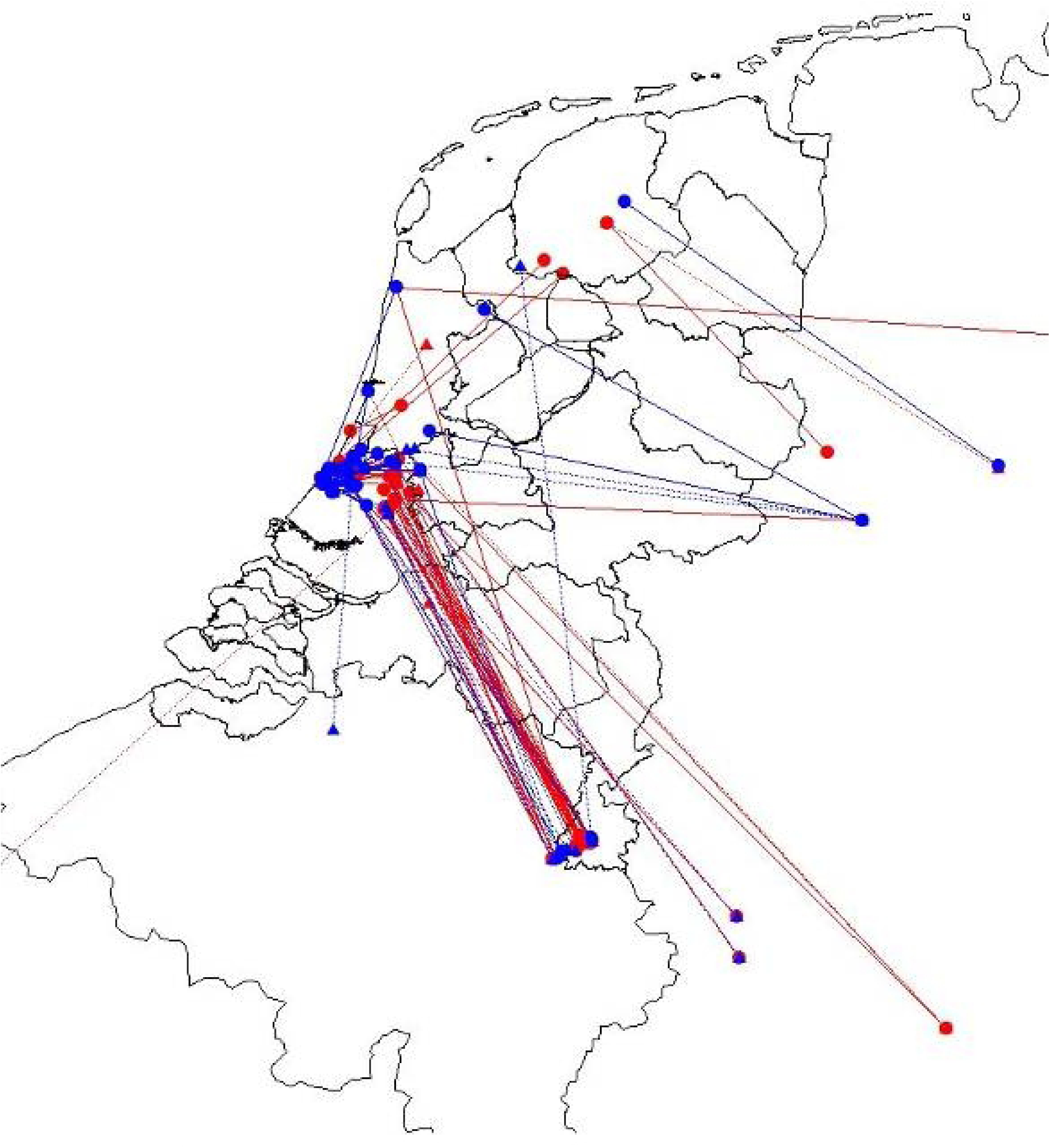
Map of recoveries from former (a) and recent (b) migration studies. In both figures males are shown in blue and females in red. The age of an animal is indicated by the shape of the points (square = unknowns age, circles = adult, triangle = sub-adult). The lines connect the point of capture and the point of recovery (adult / unknown age = unbroken line, sub-adult = broken line).

After 1980 a total of 1,727 pond bats were captured and marked (195 in their winter habitat and 1,532 in their summer habitat, Table 1a). Of the 193 recaptures, 127 (97 males and 30 females) were found in the bunkers near their summer habitat. Only a small number of animals were recovered at distances greater than 50 km from their original place of capture. These long distance migrants, both adult and sub-adult, were recovered in hibernacula such as the fortresses in Antwerp, bunkers in Calais (France), several sites in the Teutoburg forest region, limestone mines in Limburg and underground caves in the Eifel (Fig. 3b). No recoveries were made in the province of Namur (southern Belgium).

## Discussion

We find evidence for the energetic trade-off between mating and migration by looking at confounding factors such as BCI. The overall ratio of males and females present in the hibernacula investigated in this study was not uniform, we found typical male and female assemblages. The typical migration distance in the first study period (1930-1980) far exceeds that of the distance during the second study period (1980-2015). Therefore, in accordance with the results of this study, we argue that males after spending energy on mating, prefer to hibernate locally. Indeed, we find a strong increase in the population size in the local hibernacula. Taken together, these results indicate that the trade-off between hibernation, mating and migration is present.

Several other authors showed energy spend during migration has a large impact on life history. This paper leads to new insights how migration, mating and hibernation interact. Our finding that males choices led to settlement of new (before 1945 not existing) local hibernacula, followed with a change of migration patterns, are unique. Our study exemplifies how bats were able to adapt, within 40 year time, to a changed habitat. Unfavourable anthropogenic activities in foraging habitat and near the maternity dwellings are happening in a very rapid tempo. Perhaps our results can help define conservation priorities and timelines, to prevent endangered bats from going extinct.

All bats seem promiscuous, both females and males mate several times. Females tend to visit the males in their mating sites. In some species multiple maternity colonies make use of one mating site, both throughout the mating season and on individual nights [57]. In other species, such as *Myotis nattereri*, it has been shown that females from a single maternity colony attend multiple mating sites [58]. Other studies show that females visit these mating sites only for one or several nights, then returning to their maternity roost [59, 60]. During a telemetry project on pond bats, not linked to this study, the first author observed a similar pattern of alternate use of maternity and mating site. This mating system is only possible when mating sites are within short flying distance of maternity sites. While pond bats are considered long distance migrators, with the settling of the males in local hibernacula, we assume they managed to reinstall their favoured mating system.

The trade-off between mating, migration and hibernation is also seen in other mammalian clades, such as ungulates and seals [e.g. 61, 62]. In such systems males spend a lot of energy defending females and are considered territorial. Male bats have a similar mating behaviour, known as swarming [63]. Swarming behaviour, with intense flight activity, circling in and around the entrance of a site, is considered an interspecies social event and besides mating it also believed to play an important role in the assessment of the suitability of a hibernaculum [64] and/ or information transfer regarding its location [65]. Unlike other bat species, pond bats tend to spend little time swarming in front of the entrance, but spend more time outside making long patrol flights to and from the entrance. Presence of large groups of pond bats in the bunkers in Zuid-Holland in the period August –October suggest that they spend their time in temporary harems, like *Myotis myotis* [66]. Given that we do not have individualized data of the behaviour of male bats, we cannot determine the exact nature how they spend their energy. Future studies into the mating behaviour of the pond bat can help draw this conclusion.

On basis of our biometry data it can be understood that males have the largest gain in the change of hibernacula closer to the summer habitat, as their fatting up period between mating season and hibernation is shortest. Their trade-off between the risk of choosing a new, maybe not suitable hibernaculum, and a longer trip to the well-known hibernacula seems to pay off, as their numbers are increasing rapidly. Possible additional advantages are the increase in the length of the mating season (and thus greater chance of siring offspring) and a strong first male advantage. Other studies suggest the number of offspring sired is significantly affected by the position in the mating sequence [67], so this increased first male advantage may be a significant gain. The first male advantage is also known with another bat species, such as *Rhinolophus ferrumequinum* [68].

### Conservation implications and applications

Our findings have several notable implications for monitoring surveys. The non-uniform distribution of males and females across hibernacula can influence the monitoring results and lead to false conclusions. Whereas a change in population size in a female site may reflect the trend during the summer, similar changes in population size in a male biased site may only reflect a changed preference of site selection. Therefore, we advise to form conclusions about observed changes in population trend based on the hindsight of results presented in this paper.

The energetic arguments presented in this paper confirm that the function of hibernacula for a population of bats as a whole can differ; from male biased assemblages with a predominant role in mating to females biased assemblages with a predominant role for stable hibernation. In accordance to their function such sites may also differ critically in terms of their conservation value for the corresponding species. In addition, protection measurements and microclimate preferences for such site may also differ accordingly.

